# UniFed: A unified deep learning framework for segmentation of partially labelled, distributed neuroimaging data

**DOI:** 10.1101/2024.02.05.578912

**Authors:** Nicola K Dinsdale, Mark Jenkinson, Ana IL Namburete

**Affiliations:** Oxford Machine Learning in NeuroImaging (OMNI) Lab, Department of Computer Science, University of Oxford; Wellcome Centre for Integrative Neuroimaging, FMRIB, University of Oxford; Australian Institute for Machine Learning (AIML), Department of Computer Science, University of Adelaide; South Australian Health and Medical Research Institute (SAHMRI)

## Abstract

It is essential to be able to combine datasets across imaging centres to represent the breadth of biological variability present in clinical populations. This, however, leads to two challenges: an increase in non-biological variance due to scanner differences, known as the *harmonisation* problem, and, data privacy concerns due to the inherently personal nature of medical images. Federated learning has been proposed to train deep learning models on distributed data; however, the majority of approaches assume fully labelled data at each participating site, which is unlikely to exist due to the time and skill required to produce manual segmentation labels. Further, they assume all of the sites are available for training. Thus, we introduce `UniFed`, a unified federated harmonisation framework that enables three key processes to be completed: 1) the training of a federated harmonisation network, 2) the selection of the most appropriate pretrained model for a new unseen site, and 3) the incorporation of a new site into the harmonised federation. We show that when working with partially labelled distributed datasets, `UniFed`produces high-quality segmentations and enable all sites to benefit from the knowledge of the federation. The code is available at https://github.com/nkdinsdale/UniFed.

---

The convergence of deep learning (DL) and big data in neuroimaging has led to the exploration of new research questions in a data-driven manner. Fuelled by their capacity to capture intricate, non-linear relationships and patterns within data [1], DL techniques have been employed across a broad spectrum of neuroimaging applications, often achieving state-of-the-art performance, such as brain age prediction [2], segmentation [3] and registration [4]. However, the complexity and subtleties of neuroimaging data have, thus far, limited the application of DL techniques beyond the research domain [5]. Research MR images often differ substantially from those collected in clinical environments [6] – for instance, in imaging protocol and the demographic population being imaged – limiting the clinical applicability of models trained on research scans.

It is, therefore, necessary for DL models to be trained on data that are representative of the clinical populations of interest [7]. Despite the development of some large-scale neuroimaging datasets, such as the UK Biobank [8], the majority of neuroimaging datasets remain small, especially for rare clinical conditions. Thus, there is a need to pool data across sites and scanners to increase the amount of data available and the population represented by the data. This however leads to two new challenges: *harmonisation* and *data privacy*. The harmonisation problem refers to different MRI scanners and protocols creating images with different characteristics. Pooling these images leads to an increase in non-biological variance [9], masking the biological signal of interest [10]. The data privacy problem arises because MR images of the brain are inherently personal information and their sharing is protected by legislation such as HIPPA [11] and GDPR [12].

Federated learning (FL) has been proposed as a method to train models on distributed data [13]. The data remain at their local site-specific servers, rather than being shared centrally, and each site trains a local models on its own private datasets. The local sites then only share the weights or gradients of the local learned model, which are aggregated to create the global model, which is able to represent the data across the sites. FL has the potential to become the standard paradigm for multisite neuroimaging studies, with previous works demonstrating its feasibility and potential [14, 15].

However, the majority of FL approaches assume that all of the sites possess fully labelled data [16, 17] and that all of the sites are available and accessible when training the federation [18]. When working with neuroimaging data, especially that collected within a hospital environment, neither of these assumptions are likely to hold. We, therefore, propose `UniFed`, a unified federated harmonisation framework, which enables three key processes to be completed: 1) the training of a federated harmonisation network, 2) the selection of the most appropriate pretrained model for a new unseen site, and 3) the incorporation of a new site into the harmonised federation. The framework is general and could be applied to many architectures and tasks. We demonstrate it for semantic segmentation of subcortical grey matter structures, as it forms a key part of many neuroimaging analyses [6].

The three components of `UniFed`are based on the observation that the features can be encoded as Gaussian distributions. This enables us to encode and share the feature distributions for each site without sharing individual’s information: as the distributions are calculated across the population for each site, they reduce the leakage of personally identifying information.

We demonstrate that `UniFed`enables federations to be trained in low label regimes, with the sharing of feature statistics allowing the sites in the federation to benefit from the pool of knowledge across sites while maintaining individual privacy. The ability to train federated networks on partially labelled datasets is vital, especially for medical images, due to the difficulty in obtaining high quality manual labels. Manual labels created by clinicians are regarded as the gold standard but their availability is limited due to the time and expertise required to produce them. We show that our federated models generalise better to the sites that were unseen by the federation. Further, the subsequent two stages of `UniFed`enable us to maximise the performance on unseen sites, through the use of the shared feature statistics, without requiring any new sites to have manual labels available. First, the model selection approach enables us to select the best model from a *model zoo* of pretrained models, maximising the performance for the unseen site, based on choosing the model with the most similar reference feature distribution to that of our new site. Second, the model adaptation approach allows us to finetune the model for the new site, without requiring access to labels, by aligning the feature distributions. These enable new sites to benefit from the knowledge of the federation, without model retraining or needing access to the training data.

## Results

We simulated a data federation using 16 sites from the ABIDE dataset [19], considering the task of segmenting four subcortical regions of the brain (brain stem, thalamus, putamen and hippocampus) from the T1 structural images. The sites were split into four categories: *Reference Site*, a fully labelled site (NYU) is always available for training and possesses the most labelled data; *Labelled Sites in Federation* (0-5 sites); *Unlabelled Sites in Federation* (5-10 sites), and *Unseen Sites* (5 sites). We explored within these splits a range of data scenarios. Our results show that `UniFed`is able to produce high quality segmentation results across the data scenarios considered, while not requiring the sharing of personal information, showing its potential to enable the training of high-quality DL models for partially labelled (semi-supervised) distributed data.

### Federated Learning Framework for Partially Labelled Datasets

We considered two ways in which the sites within the federation can be partially labelled: some sites being fully labelled and some unlabelled, and all supervised sites being partially labelled.

#### Increasing Number of Supervised Sites

The case where labels only exist for a few sites in the federation is potentially the most challenging for non IID data, such as data with varying scanner characteristics across sites.

Table 1 shows the results when increasing numbers of sites (1 (only the reference site) to 6) within the federation are fully labelled, with Dice scores averaged across subcortical regions and across the sites, separately for the sites within the federation and the unseen sites. We compared `UniFed`to a range of federated methods, including domain adaptation approaches (FADA [20] and FedHarmony [21]) and semi-supervised FL methods (FedPerl [22] and FSSL-DPL [23]). For the case where only the reference site was labelled (NYU), the domain adaptation approaches and `UniFed`showed similar performance (*>* 66% Dice), with no significant difference between the results for the best method and `UniFed`. For the remaining cases (2-6 sites labelled) `UniFed`outperformed the existing methods. A limited subset of these experiments were completed for a different random split of the sites and a different choice of reference site, and the same trend was seen (available in the supplementary material).

**Table 1:**
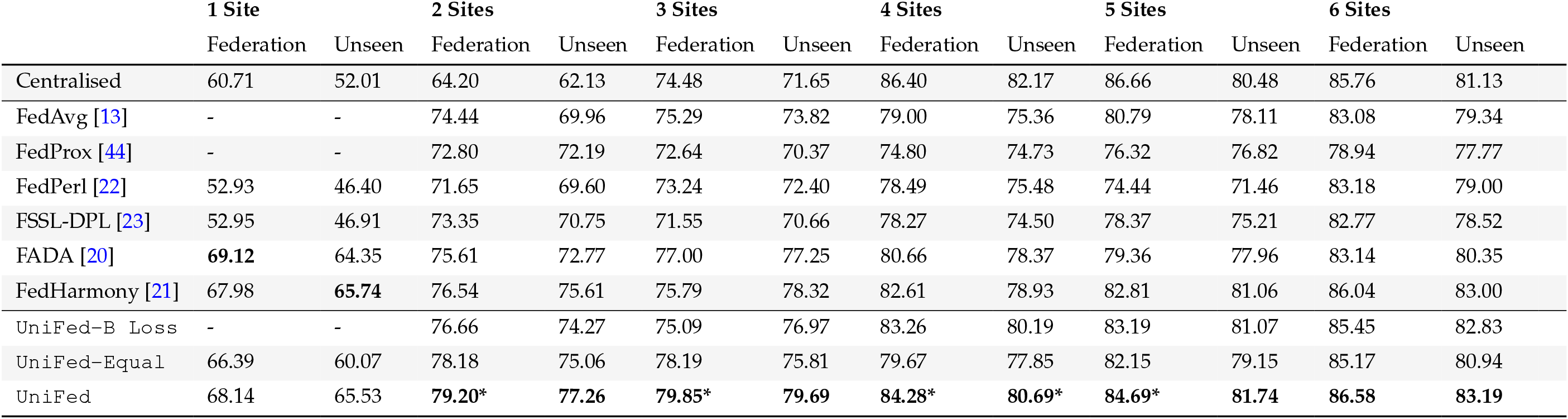
Partially Labelled FL Results - Sites Labelled. The results for the partially labelled FL framework are presented, for increasing numbers of fully supervised sites within the federation. The Dice score is averaged over the 4 tissue classes. `Federation`are the Dice values averaged across the tissue classes, for sites within the federation, and `Unseen`are the Dice values averaged across the unseen sites. `UniFed - B loss`= ablation study removing the Bhattacharyya loss, `UniFed - Equal`= ablation study using FedAvg as the default aggregation scheme. The best performing method is in bold. ^*^ indicates statistical significance between best method and next performing method (p<0.01).

|  | 1 Site |  |  | 2 Sites |  |  | 3 Sites |  |  | 4 Sites |  |  | 5 Sites |  |  | 6 Sites |  |
| --- | --- | --- | --- | --- | --- | --- | --- | --- | --- | --- | --- | --- | --- | --- | --- | --- | --- |
|  | Federation | Unseen |  | Federation | Unseen |  | Federation | Unseen |  | Federation | Unseen |  | Federation | Unseen |  | Federation | Unseen |
| Centralised | 60.71 | 52.01 |  | 64.20 | 62.13 |  | 74.48 | 71.65 |  | 86.40 | 82.17 |  | 86.66 | 80.48 |  | 85.76 | 81.13 |
| FedAvg [13] | - | - |  | 74.44 | 69.96 |  | 75.29 | 73.82 |  | 79.00 | 75.36 |  | 80.79 | 78.11 |  | 83.08 | 79.34 |
| FedProx [44] | - | - |  | 72.80 | 72.19 |  | 72.64 | 70.37 |  | 74.80 | 74.73 |  | 76.32 | 76.82 |  | 78.94 | 77.77 |
| FedPerl [22] | 52.93 | 46.40 |  | 71.65 | 69.60 |  | 73.24 | 72.40 |  | 78.49 | 75.48 |  | 74.44 | 71.46 |  | 83.18 | 79.00 |
| FSSL-DPL [23] | 52.95 | 46.91 |  | 73.35 | 70.75 |  | 71.55 | 70.66 |  | 78.27 | 74.50 |  | 78.37 | 75.21 |  | 82.77 | 78.52 |
| FADA [20] | **69.12** | 64.35 |  | 75.61 | 72.77 |  | 77.00 | 77.25 |  | 80.66 | 78.37 |  | 79.36 | 77.96 |  | 83.14 | 80.35 |
| FedHarmony [21] | 67.98 | **65.74** |  | 76.54 | 75.61 |  | 75.79 | 78.32 |  | 82.61 | 78.93 |  | 82.81 | 81.06 |  | 86.04 | 83.00 |
| UniFed-B<sub>Loss</sub> | - | - |  | 76.66 | 74.27 |  | 75.09 | 76.97 |  | 83.26 | 80.19 |  | 83.19 | 81.07 |  | 85.45 | 82.83 |
| UniFed-Equal | 66.39 | 60.07 |  | 78.18 | 75.06 |  | 78.19 | 75.81 |  | 79.67 | 77.85 |  | 82.15 | 79.15 |  | 85.17 | 80.94 |
| UniFed | 68.14 | 65.53 |  | **79.20\*** | **77.26** |  | **79.85\*** | **79.69** |  | **84.28\*** | **80.69\*** |  | **84.69\*** | **81.74** |  | **86.58** | **83.19** |

We also conducted an ablation study, removing both of our additions to the vanilla FedAvg training procedure: the Bhattacharyya loss (`UniFed - B Loss`(eq. 20)) and the equal aggregation (eq. 14), replacing it with the FedAvg aggregation scheme (`UniFed - Equal`). Removing both components leads to a decrease (*>* 1% Dice) in performance compared to `UniFed`, showing that both components are integral to the performance of `UniFed`in this setting.

Figure 2 provides further quantitative and qualitative results for the federated results where two sites are fully labelled: the reference site (NYU) and, additionally, Leuven. Figure 2a) shows the boxplots for FedAvg, FADA and `UniFed`broken down by site. As would be expected, the performance is better for the labelled sites (NYU and Leuven) than for most of the unlabelled sites, but it is evident that `UniFed`leads to an improvement across all of the sites, including the unseen sites. Figure 2b) shows qualitative results for a labelled and unlabelled site from the federation: Leuven (labelled) and KKI (unlabelled), comparing the single-site source model (NYU only), FedAvg, FADA, and `UniFed`. Axial, sagittal and coronal views are shown centred around the regions of interest. Finally, figure 2c) shows representative features shared for the FADA method and for `UniFed`. For FADA the features shown are a single slice of the 3D feature arrays, where eight 3D feature arrays per subject, per site are shared. For `UniFed`, the features shown represent all the shared information for a given site (reshaped from a vector for ease of presentation).

**Figure 1.**
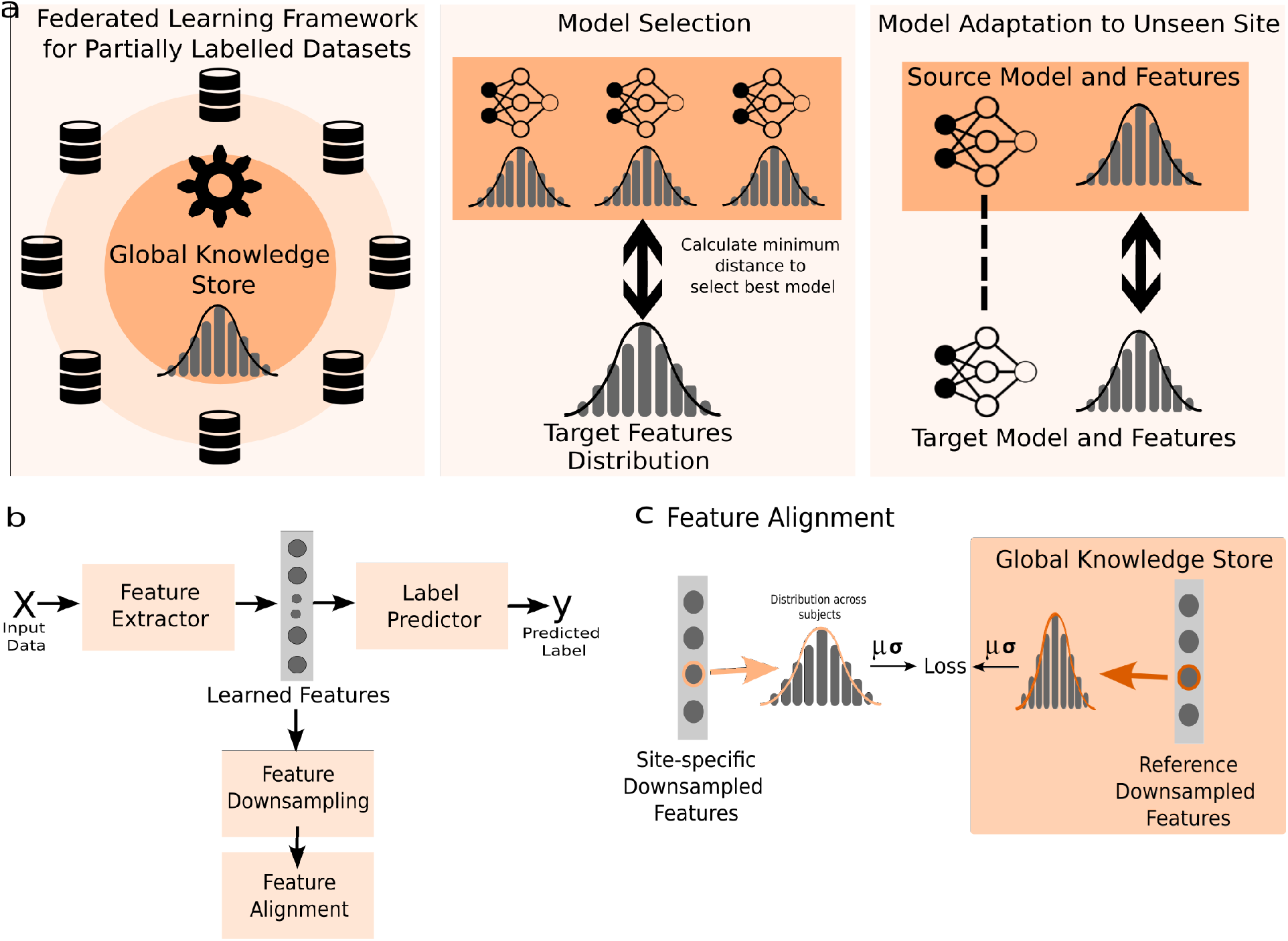
Overall UniFedframework. The framework is formed of three components, which are shown in (a). The first is a FL framework, which enables training on distributed datasets. The second component is model selection, which for a new target dataset enables the best model for that target data to be selected from the ‘*model zoo*’: the collection of available models. The final component is model adaptation, which allows the model to be updated to improve the performance on an unlabelled target dataset. The three components are based around the core idea of modelling features as Gaussian distributions across populations, enabling model adaptation with limited communication. In b) the model framework is shown, where the form of the feature extractor and label predictor depend on the task of interest and the study preferences. Here, we use a 3D UNet. The learned features are taken from the penultimate convolutional block, and are then downsampled before the feature alignment can be completed (c). For feature alignment, each feature is modelled as a 1D Gaussian distribution and then compared to the global distribution across the sites by comparing the distribution statistics - the mean (*µ*) and standard deviation (*σ*). b) and c) are used across all three stages of the framework.

**Figure 2.**
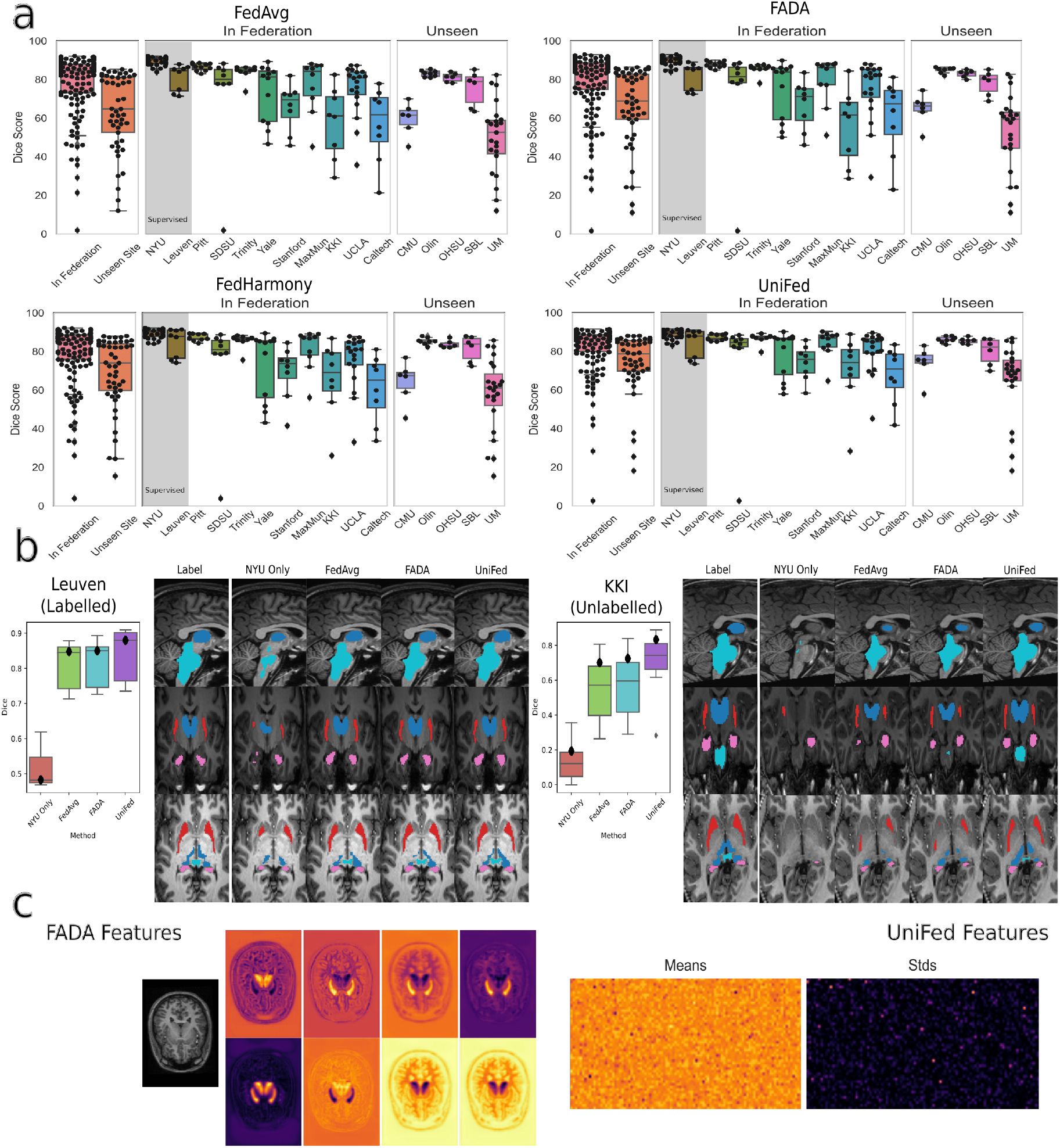
Quantitative and qualitative results for the partially labelled federated setting with 2 sites supervised. (a) Boxplots of the dice scores for FedAvg, FADA and `UniFed`, showing the performance of the federation and unseen sites, and broken down by site. Points show performance for each test subject. (b) Qualitative segmentation results for a labelled site (Leuven) and unlabelled site (KKI) showing an axial, sagittal and coronal view, cropped around the region of interest. Light blue = brain stem, Dark blue = thalamus, pink = hippocampus, red = putamen. c) Examples of the features shared by FADA and `UniFed`(reshaped from a vector for ease of presentation). For FADA these are the features from a single subject, and for `UniFed`they represent the shared statistics for a whole site.

#### Increasing Percentage of Supervision

We then considered the other way in which the sites could be partially labelled: when the *Labelled Sites in Federation* all have a small percentage of labelled subjects. Thus, we tested having the fully supervised reference site (NYU) and increasing percentages of the images for the five further labelled sites in the federation. We compared to domain adaptation approaches (FADA [20] and FedHarmony [21]) and also to a locally semi-supervised method (omnisupervised learning following [24]). It can be seen in Table 2 that, for all non-zero percentages of labelled data considered, `UniFed`outperformed the existing methods. The improvement was greater when a low percentage of labelled data was considered. As before, we also completed an ablation study, again demonstrating the benefits of both components to `UniFed`. Most likely due to the extreme imbalance in the number of samples at the different sites, the benefit of aggregating equally across all of the sites is especially clear.

**Table 2:** Partially Labelled FL Results - % Labelled. The results for the partially labelled FL framework are presented, when we have the fully supervised reference site and an increasing percentage of labelled data at each of the remaining five labelled sites in the federation. The Dice score is presented averaged over the four tissue classes. `Federation`are the Dice values averaged across the sites from within the federation, and `Unseen`are the Dice values averaged across the unseen sites. The best performing method is in bold, star indicates statistical significance between best method and next performing method (p<0.01).

|  | 0% |  | 1% |  | 5% |  | 10% |  | 20% |  |
| --- | --- | --- | --- | --- | --- | --- | --- | --- | --- | --- |
|  | Federation | Unseen | Federation | Unseen | Federation | Unseen | Federation | Unseen | Federation | Unseen |
| Centralised | 60.71 | 52.01 | 63.44 | 60.10 | 82.45 | 80.04 | 85.15 | 80.36 | 85.29 | 80.50 |
| FedAvg [13] | - | - | 68.11 | 64.65 | 67.79 | 60.37 | 69.70 | 65.28 | 72.67 | 67.97 |
| Omnisupervised [24] | - | - | 61.84 | 55.35 | 62.45 | 58.68 | 61.99 | 57.22 | 65.15 | 60.40 |
| FedProx [44] | - | - | 69.41 | 69.08 | 69.64 | 63.75 | 71.17 | 69.90 | 71.76 | 69.76 |
| FADA [20] | **69.12** | 64.35 | 71.74 | 66.71 | 72.39 | 68.06 | 71.97 | 67.66 | 70.09 | 66.06 |
| FedHarmony [21] | 67.98 | **65.74** | 71.21 | 73.74 | 79.06 | 77.69 | 81.54 | 79.16 | 83.89 | 81.19 |
| UniFed - B Loss | - | - | 70.25 | 73.33 | 78.81 | 77.55 | 79.66 | 76.62 | 81.25 | 77.04 |
| UniFed - Equal | 66.39 | 60.07 | 68.72 | 63.48 | 68.44 | 61.89 | 69.68 | 64.20 | 75.38 | 67.80 |
| UniFed | 68.14 | 65.53 | **76.46\*** | **74.97\*** | **81.81\*** | **78.59\*** | **82.01\*** | **79.33** | **84.18** | **81.94** |

### Model Selection

We wish to select the best from a collection of pretrained models (the *model zoo*) for our new unseen site, in the absence of target labels. For our model zoo, we utilise the models trained for the experiments in Table 1, and so have potential source models from across the different comparison methods and number of labelled sites involved in training. Figure 3 shows the Dice scores plotted against the Bhattacharyya distances between features for the different pretrained models, for each unseen site separately. For each unseen site there is a negative correlation between the Bhattacharyya distance and the average Dice score (UM: r= -0.850 p= 6.94*×*10^*−*12^, CMU: r= -0.461 p= 0.003, OHSU: r= -0.206 p= 0.021, SBL: r= -0.465 p= 0.003, Olin: r= -0.467 p= 0.003), showing that there is a correlation between Dice score and Bhattacharyya distance, and that the Bhattacharyya distance between the reference features and target features can be used for model selection. The `UniFed`trained models were selected for 4 of the 5 unseen sites, showing that the domain adaptation creates features that generalise better, even for unseen sites. As would be expected, the models trained with the largest number of labelled sites generalise best to the unseen sites.

**Figure 3.**
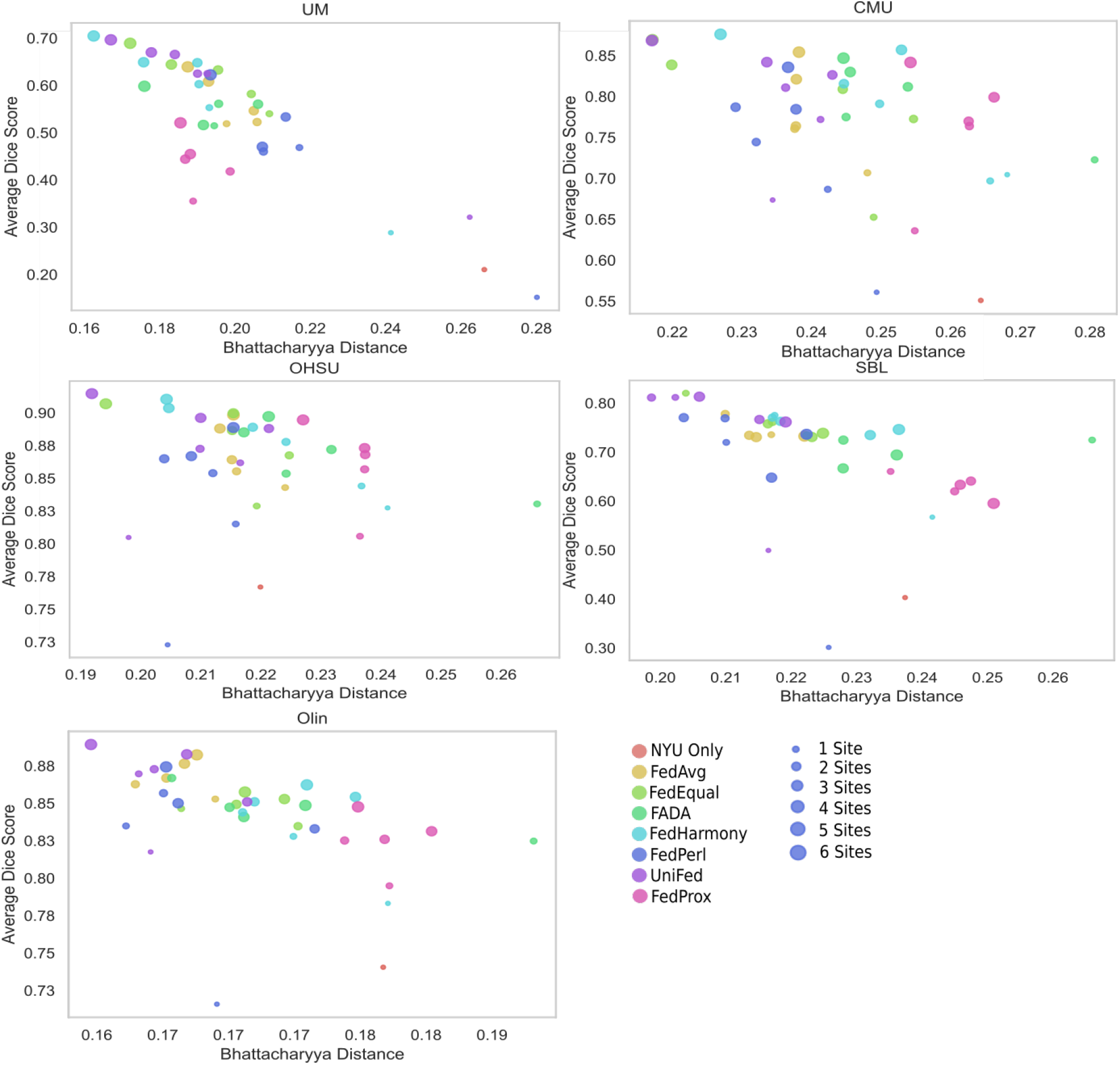
Best Model Selection Results. For each unseen site, the Dice score produced by a model from the model zoo is plotted against the Bhattacharyya distance between the reference feature distributions and the unseen site feature distribution. The models trained for Table 1 were used as the model zoo, across the different methods and number of sites used in training. The colour of the marker corresponds to the method used to train the model and the size of the marker corresponds to the number of labelled sites that were used to train the model. Dice score clearly correlated with the Bhattacharyya distance and the shorter the distance, the more similar the feature distributions, meaning that the Bhattacharyya distance can be used for model selection.

### Model Adaptation to Unseen Site

To explore adapting the model to an unseen site, we considered two source models - the NYU only trained source model and the `UniFed`trained model for all six sites labelled. Each source model was adapted to all of the five unseen sites, and the reported result is the Dice score averaged across all four subcortical regions, and across all of the unseen sites. In Table 3 it can clearly be seen that the NYU only source model had much lower performance on the unseen sites compared to the `UniFed`model trained on 6 labelled sites (52.01 vs 83.14). Although adapting the NYU only source model led to a greater increase in performance (52.01 to 68.71), the performance was still than the `UniFed`model trained on 6 labelled sites. We compared `UniFed`to existing source-free domain adaptation methods for segmentation (ADAEnt [25] and ADAMI [26]), which performed badly due to the extreme label imbalance and small batchsize. We also compared to minimising the MMD between the features. `UniFed`outperformed the existing methods for both source models. `UniFed-EM`shows an ablation experiment in which the expectation maximisation to fit the Gaussian distribution is removed. This component provides memory between batches, and its removal leads to a substantial drop in performance.

**Table 3:** Model Adaptation Results. The results are shown for adapting a source model to the unseen sites, with the Dice score averaged over the unseen sites. Two source models were considered, the NYU only source model and the `UniFed`model trained with all six sites supervised. The best method is shown in bold.

|  | NYU Only Source Model | UniFed 6 Sites Source Model |
| --- | --- | --- |
| Source Model | 52.01 ± 21.55 | 83.19 ± 7.09 |
| Target Finetune (Fully Labelled Oracle) | 85.50 ± 4.21 | 87.49 ± 3.99 |
| Entropy | 0.166 ± 0.109 | 4.54 ± 2.83 |
| ADAEnt [25] | 0.780 ± 0.978 | 10.57 ± 8.45 |
| ADAMI [26] | 0.292 ± 0.367 | 8.83 ± 11.61 |
| MMD | 66.79 ± 12.46 | 84.37 ± 6.02 |
| UniFed - EM | 60.45 ± 15.88 | 84.09 ± 6.20 |
| UniFed | **68.71 ± 12.52** | **85.43 ± 6.13** |

## Discussion

Medical images are inherently personal and their sharing is protected by data privacy laws. Thus, the development of methods that work with distributed data is crucial, and increasingly important for the creation of DL methods appropriate for clinical application. We have demonstrated the three aspects of our unified framework that tackle three major tasks when working with distributed data: FL for partially-labelled datasets, model selection, and model adaptation to a new, unseen site.

Our framework requires only the sharing of the summary statistics (the mean and standard deviations of the Gaussian distributions), not the underlying features or data. The benefit of this can clearly be seen in Fig. 2c) that shows the shared features for FADA and the summary statistics for `UniFed`. First, the features are much larger in size (height x width x depth x channels x number of subjects) compared to the summary statistics (subsampled features (1000) x 2 – mean and standard deviation). Second, the features contain a large amount of the information contained in the original images and, thus, sharing them would likely leak personal information. It also enables us to use the Bhattacharrya distance in all three stages: the Bhattacharyya distance measures the distance between two distributions, with a simple closed form solution for Gaussian distributions, making it efficient and simple to calculate.

The first step of `UniFed`(federated learning with partially labelled datasets) matches the performance of existing approaches for partially labelled FL or outperforms them. The vast majority of existing methods for training federated models assume either fully supervised data at the local site [16, 17], or make assumptions specific to classification tasks, meaning that they cannot be readily applied to segmentation tasks [27]. Further, for a site to benefit from the knowledge within the federation, it is normally required for the site to be available at training time [18]. Existing methods are often adversarial [20, 21], and thus require the training of a discriminator, which presents a challenge for the training and balancing of multiple losses and learning rates [28], whereas our method only has a single additional loss function, lowering the barrier to applying the approach to different tasks and models. The only additional hyperparameter introduced is the weight between the main task loss and the Bhattacharyya loss (*λ*, Eq. 13), and we found that the result was consistent across a wide range of values (see Supplementary Material). Other approaches are based on using the local models for the labelled sites to create pseudo labels for the unsupervised sites [22, 23]. However, for our dataset there is a large amount of domain shift between the different local sites and the task is relatively challenging; thus, the pseudo labels created are often of too low quality to produce acceptable segmentations, especially when few sites have labels available. This task is vital as the time and expertise required to create manual segmentation labels for neuroimaging datasets means that it is very rare to have fully labelled datasets available at all local sites.

Furthermore, `UniFed`learns a more generalisable feature representation, leading to a large improvement in performance on the entirely *unseen sites* compared to all of the other methods considered. The adversarial approaches and `UniFed`are both able to consider the data distributions of the unlabelled datasets within the federation, giving an improvement in performance on these sites. This can be seen in Fig 2a) by comparing the federation results from FedAvg to FADA, FedHarmony and `UniFed`with the greatest improvement obtained when using `UniFed`. The improvement can also be seen for the Unseen sites, showing that `UniFed`learns features that generalise well beyond sites available during training. Qualitative results (Fig. 2b) also show clearly the improvement gained by using `UniFed`. For instance, for the KKI segmentations, the degree of the domain shift between the NYU training data and the KKI site is clear: the *NYU only source model* entirely fails to segment the majority of the structures, clearly indicating the need for our approach, as the base segmentations would be entirely unusable. Qualitatively, it is evident that `UniFed`produced the best segmentation prediction, across the four structures.

The degree of domain shift observed explains the poor performance [29] of the pseudo label-based approaches [22, 23] and also the poor performance of the omnisupervised-based approach [24] for the case when a percentage of data points were labelled at each site (Table 2). For the percentage supervised experiments, `UniFed`again consistently outperforms the existing methods. The relatively poor performance of FADA – despite the sharing of the whole features and the ablation study (`UniFed - Equal`) – show the importance of the equal aggregation step (as per Eq. 14) especially when the number of samples is very imbalanced. Thus, our proposed approach is applicable to the two forms of partially labelled local sites, with no changes to the approach.

The second part of the framework is model selection: selecting the best model to use for a new dataset is vital when limited or no labels are available for the new site. This is clearly challenging as we cannot simply select the best model based on performance for the desired metric. Although some methods exist for predicting model performance [30, 31], which could then be used to select the best model for the new site, these mostly consider classification. We hypothesised that the Bhattacharyya distance between the reference and the new model features could be used to select the best site: the shorter the distance, the more similar the feature representations. As we are already sharing the feature distributions in our framework, this can be assessed for free without requiring any additional information to be shared. For each unseen site, there was a clear negative correlation between the Bhattacharyya distance and the average Dice score, showing that the Bhattacharyya distance is a good metric to use for model selection. Although the model selected is not always strictly the highest Dice performance, the selected model consistently performs highly and there is no significant difference between the best model and the selected model for any of the unseen sites (p<0.05).

The importance of the model selection becomes clear when we consider the final part of the framework: model adaptation. Despite the improvement in performance for adapting the NYU only source model, the performance was still worse than the `UniFed`trained model, showing that, although model adaptation can greatly increase the performance of the model without any need for labels, it cannot fully account for the selection of a bad source model in the first place.

Model adaptation is an important part of the framework as it allows us to create models which are suitable for new, unlabelled datasets. Our approach provides the best performance across the unseen sites, while only requiring the storing and sharing of a single set of aggregate mean and standard deviations in addition to the model weights. We treat the problem as a s*ource-free domain adaptation* problem, which means we can adapt the model to the new site without needing access to the other sites in the federation. This is important for two reasons: first, retraining the federated model with the new site is expensive in terms of both time and resources; second, it allows the model to be adapted when the federated data are not accessible – for instance, if the new site were not part of the data-sharing agreement or if the data were no longer available. The majority of existing approaches are either generative, which are inappropriate for medical images due to the difficulty of training and the likelihood of data leakage [32], or rely on the minimisation of the model entropy, based on the fact that more confident predictions are more likely to be correct. We found that the entropy-based methods collapsed for our task, likely due to the extreme class imbalance – the majority of the volumes are occupied by background – and the small batchsize that can be achieved when training models with 3D volumes.

Although our framework has demonstrated strong performance across the three challenges, our study is not without limitations. First, we have considered only a single, albeit multisite, dataset and task. The ABIDE dataset [19] was selected due to the high number of sites within the federation, and the task of subcortical segmentation due to the relative difficulty of the task. We expect that the results would generalise to other datasets and we demonstrated the applicability of modelling the features as a Gaussian distribution for many tasks and datasets in our previous work [33]. The framework is general, with no dataset-specific assumptions, and so we expect the results to generalise well.

In our work, we have made the assumption that a single fully-labelled reference site is available. This is a reasonably likely scenario – for instance, where a site is leading the study – but it does present a potential limitation. It is, however, possibly unnecessary so long as sufficient labelled samples are available across the federation to learn a high performing model. Our results were robust to the choice of reference site (see Supplementary Material).

Finally, our method assumes that there are sufficient data available for a given site, whether for training within the federation or for model adaptation. The individual sites in the ABIDE dataset are small (smallest dataset size was 27 subjects after preprocessing), and we were able to train the networks using only basic augmentation, but there will be a limit to the smallest site usable. Exploration of test time adaptation methods [34, 35] would possibly be the best approach for very small dataset sizes, as they can enable model adaptation with a single sample.

We have presented `UniFed`, a unified federated harmonisation framework that enables the training of high performing models for distributed and partially labelled datasets. The three parts: training of a federated network, model selection and model adaptation, are linked by the simple modelling of the features as a Gaussian distribution. Therefore, the approach is general and widely applicable to different segmentation tasks and choices of model architecture, and thus, to many distributed imaging studies.

## Methods

The code is available at https://github.com/nkdinsdale/UniFed.

### Data

To simulate a multisite federation, we utilise the ABIDE data [19], with the imaging data acquired from each site being held in a separate data silo for that given site. Thus, each site is regarded as a separate domain, following [36]. The T1 images were preprocessed using the FSL Anatomical Processing Script^1^ (`FSL_anat`), which includes orientation to MNI space, bias field correction and linear registration to the MNI template. Only subjects that completed the whole pipeline were kept. This led to the largest individual site (NYU) having a total of 181 subjects available and the smallest site (CMU) having only 27 subjects in total. The preprocessing pipeline failed to complete for almost all of the data from one site (USM), critically including the generation of segmentation labels used for model training, and thus was entirely discarded.

For the segmentation labels, proxy labels produced by FSL FIRST [37] (as part of the `FSL_anat`pipeline) were used. We considered segmenting four subcortical structures (both left and right combined, where appropriate): Thalamus, Putamen, Hippocampus and the Brain Stem. Only four structures were considered due to memory constraints, but were sufficient to thoroughly demonstrate the approach. All images and labels were resized to 128 *×* 240 *×* 160.

After the preprocessing, we had data from 16 separate imaging sites. We split these into four sections:

- *Reference site* - a singular site assumed to always be fully labelled. NYU was used as the reference site as it had the largest number of images after the preprocessing pipeline.
- *Labelled Sites in Federation* - sites available for training. These were randomly selected from the remaining sites: Leuven, Pitt, SDSU, Trinity and Yale.
- *Unlabelled Sites in Federation* - sites available for training but with no labels available. These were randomly selected from the remaining sites: Stanford, MaxMun, KKI, UCLA and Caltech.
- *Unseen Sites* - sites not available for training, used to assess the generalisability of the learned model. These were the remaining sites: CMU, Olin, OHSU, SBL and UM.

For each site the data were split 80%/20% into training and testing sets. All models were trained with three-fold cross validation and the result reported is for the test set, with the value averaged over the four subcortical regions.

### Network Architecture

To achieve image segmentation, we utilised a 3D UNet architecture. Details of the architecture used can be found in the Supplementary Material. We choose to share information about the distributions of the model features in the penultimate layer of the network: the point at which at the maximal amount of information has been learned from the input images and encoded by the features. If we complete domain adaptation before the final skip connection, the domain information can entirely be reincorporated [36].

### General Setup

The framework is unified by two key concepts: (i) the encoding of image features as Gaussian distributions to enable the communication of information across sites without violating data privacy, and (ii) the use of the Bhattacharyya distance to calculate the distance between sites in feature space. Using these two concepts we formulate a *Federated Learning Framework for Partially Labelled Datasets*, a *Model Selection* approach, and *Model Adaptation to Unseen Site* approach to incorporate a new site.

Although the results presented here are for segmentation, the approach is general and applicable across a wide range of tasks and model architectures. Thus, we consider a general model setup, as shown in Fig. 1c, following the framework presented by [38], where the overall network is formed of a feature extractor, with parameters Θ_*repr*_, and a label predictor, with parameters Θ_*p*_. The same network architecture design is used across all participating sites.

For a given imaging site, or domain, we have a dataset *D*^*s*^ = *{****X***^*s*^, ***Y*** ^*s*^*}* which contains T1-weighted magnetic resonance images ***X***^*s*^ and, if supervision is available for that site, also contains segmentation labels ***Y*** ^*s*^. As the sites represent data collected on different scanners, there is a domain shift present and so a model trained on any one of the sites would suffer a degradation in performance when applied to the other sites. This domain shift is explored in detail in the Supplementary Material. We thus have a multisite, partially labelled federated domain adaptation problem, where the goal is to create a single model that performs equally well across all sites.

### Federated Learning Framework for Partially Labelled Datasets

The first component of the unified domain adaptation `UniFed`is a federated learning framework, which enables the training of models across multiple sites contained in data silos. We consider applying the framework to two forms of partially labelled datasets (utilising the *Labelled Sites in Federation* subset): when increasing number of sites that have supervision available, and when the 5 partially labelled sites in the federation have increasing amounts of supervision available. Sites without supervision still contribute to the model training through feature alignment (see step 4).

**Step 1: Model Initialisation** Training is completed through multiple rounds of local training and weight communication and aggregation. First, all local models must be initialised with the same weight initialisation, Θ_*repr,t*_ and Θ_*p,t*_, for the feature extractor and label predictor respectively; *t* indicates the current weights for iteration *t*. For the first communication round, the models were all initialised with a model trained only on the NYU *Reference Site*, which is fully labelled and so could be trained as standard with full supervision. For subsequent communication rounds, the local models are initialised with the aggregated model from the previous communication round (step 5).

**Step 2: Global Information Store** The domain adaptation requires alignment of the learned feature embeddings ***Q***^*s*^ = *g*(***X***^*s*^, Θ_*repr*_) for all sites within the federation, (*Reference site, Labelled Sites in Federation* and *Unlabelled Sites in Federation*), where *s* is the current local site and *g* is the function learned by the feature extractor. To achieve this, we follow the precedent of existing privacy preserving medical imaging approaches [21, 39, 40] and create a global knowledge store to share summary statistics of the features. Previous federated learning approaches have shared the whole set of features [20]; however, for segmentation, these features will be very large and have high structural similarity with the input images. Thus, sharing the features is likely to cause data leakage. We follow [21], encoding the features as Gaussian distributions using expectation maximisation (EM), and thus the statistics to be shared are the mean and standard deviation per feature. If a Gaussian distribution were a poor fit for the features this could be expanded to a GMM, (details in the Supplementary Material). EM provides memory between batches, limiting the effect of the small batchsizes possible when performing 3D segmentation. For each feature at a local site, we fit an independent 1D Gaussian to the features, thus for feature 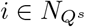 where 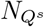 is the number of features in ***Q***^*s*^:

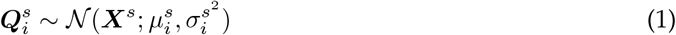

where 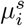 is the mean value for the *i*^*th*^ feature in site *s*, and similarly 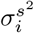 is the standard deviation for the *i*^*th*^ feature in site *s*. Thus, we have two parameter arrays that define the modelled distribution of the features, which are stored in a global information store:

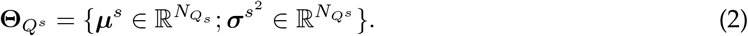

Due to representing aggregate statistics across a population, these features contain no individually identifying information. However, given how large these features are, and the high degree of redundancy between the features due to the nature of the convolutional filters, we randomly subsample the shared statistics:

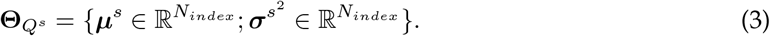

where *N*_*index*_ is the number of features retained. To be able to complete DA, the same features in the same order must be shared by each local site, and so the indices for the retained features must also be shared between sites, which clearly does not violate data privacy. Throughout our experiments, the total number of features, 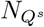, was 39,321,600, and we set *N*_*index*_ to 5000, based on the result in [33].

**Step 3: Aggregation of local distributions** The global information store contains the statistics for the local sites which we now need to aggregate to create the global reference statistics. We aggregate the statistics for all sites with supervision (*S*_*sup*_), weighted by the number of labelled image samples at each local site, *N*_*s*_, used to create the statistics:

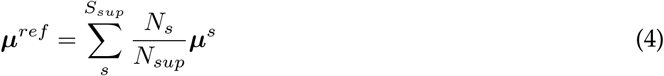

and

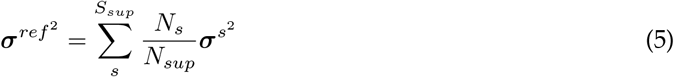

where *N*_*sup*_ is the total number of labelled images in the federation. These now form the reference features, and the following training stage aims to align the local features to these global reference features:

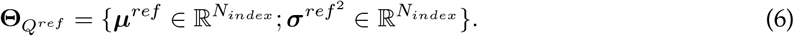

**Step 4: Training local models** The local models can now be trained for all sites in the federation. Local training is governed by two loss functions: the main task loss and the alignment loss. The main task loss can only be evaluated for samples that have labels available, forming a dataset 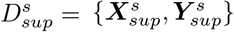

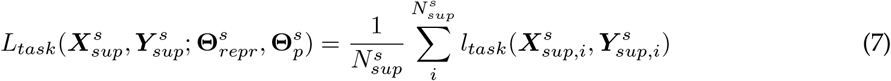

where we use Dice loss for *l*_*task*_ and 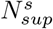 is the number of labelled samples for site *s*.

The alignment loss can be evaluated for all subjects, meaning that all subjects can contribute to the domain adaptation, including those at sites where no labels are available. The goal is to align the local source features to the global reference features. In adversarial approaches, a discriminator is added to the overall architecture to make domain invariant features. We could utilise this approach, following [21], drawing feature samples from the normal distributions; however, adversarial approaches are notoriously unstable and difficult to train. We instead align the features using the parameters of the Gaussian directly. We first fit the normal distribution to the local site, *s*, only for the subset of retained features *N*_*index*_ for all local data points *N*_*s*_:

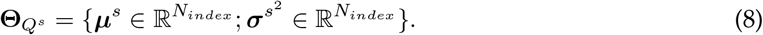

We then utilise the Bhattacharyya distance as the loss function. The Bhattacharyya distance measures the similarity of two probability distributions, which for continuous probability distributions is defined as:

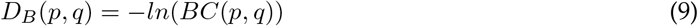

where

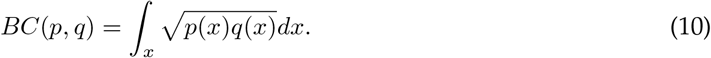

Given that both distributions are Gaussian, the Bhattacharyya distance has a simple closed form solution. If 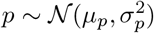 and 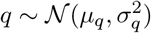 then:

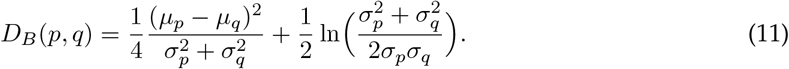

Thus, we evaluate this between our reference and local features as our alignment loss function:

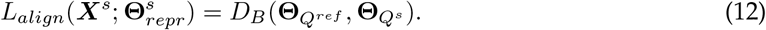

Therefore, the overall loss function is given by:

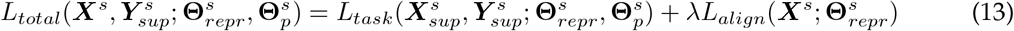

where 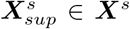 and *λ* is the relative weight between the two losses, set to 0.1 across our experiments. Clearly, for the sites which are fully unsupervised, *L*_*task*_ is zero and the training only involves the alignment of the feature distributions. The local models are all trained until they reach convergence, and patience was set to 10 epochs.

**Step 5: Model Aggregation** Once all local models have been trained to convergence, the local models are sent to the central server for aggregation to a single global model. Most aggregation schemes could be used here, but we propose to aggregate the local models equally, as in [21], due to the possible very large differences in samples available at the different sites. In this case, aggregation weighted by the number of examples, such as FedAvg [13] is likely to lead to models that specialise to the largest site, and thus still suffer a drop in performance when applied to the other sites. Thus, we aggregate the model parameters as:

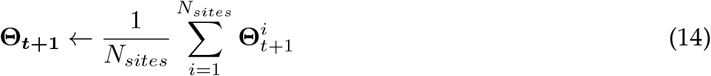

where the aggregation is completed separately for the weights of the feature extractor, Θ_*repr*_, and the label predictor, Θ_*p*_, *N*_*sites*_ is the number of sites within the federation, 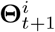 are the updated local weights, Θ_*t*+1_ are the updated global weights, and *t* is the current iteration.

The five steps are iterated through until the system reaches global convergence, leading to a final global model.

### Model Selection

Model selection assumes that a number of models have been trained by the sites in the federation. For instance, by varying multiple random initialisation seeds, different parameter values or different approaches, we form a *model zoo*. Thus, given a new, unlabelled dataset, the challenge is to select the most appropriate model for this new data. This is especially vital for source-free domain adaptation methods, which we will use to incorporate a new site into the federation, assume access to a well trained source model [41] and, thus, it is important to select an appropriate model for the unseen site. Thus we assume we have a selection of pretrained source models and a target site, *D*_*target*_ = *{****X***_*target*_*}*, for which we wish to select the best model. We hypothesise that the shorter the Bhattacharyya distance between the source and target features, the more appropriate the model will be for the target dataset. Thus, each site will have to share their feature summary statistics and model weights, but no data would need to be shared. These would be stored in the global knowledge store, and so storage of the data used for the pretrained source models would no longer be required.

At the end of the model training, for each model in the model zoo, we would need the final feature summary statistics to be calculated, aggregated across the sites that contributed to the model training. However, the features of different models may exist at different scales [42, 43], and thus the calculated distances would also be at different scales. Thus, we normalise the features for the *reference site*, NYU, using the L2 norm prior to calculating the summary statistics, such that for each feature element *i* in ***Q***^***s***^:

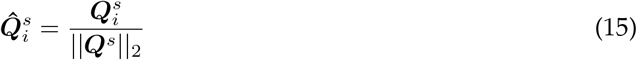

and we share the denominator as a normalisation factor for the features at all other supervised sites used in training the model. We can then calculate the aggregated summary statistics for the given reference model following Equations 4 and 5 to create the reference statistics:

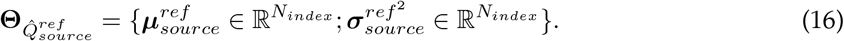

This only needs to be completed once, when the models are trained, and then stored in the global information store, until a future time when model selection is required.

When we wish to complete model selection for our new unlabelled, unseen target site, for each model in the model zoo, we calculate the statistics across all of the samples for the target site, having normalised the features using the shared normalisation factor.

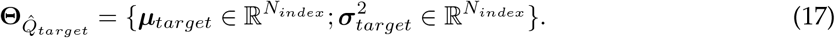

Then for each model, we calculate the Bhattacharyya distance and select the most appropriate model as the one with the shortest distance:

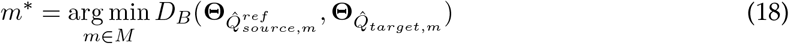

where *M* is the model zoo, and 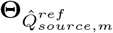 are the statistics for the *m*^*th*^ model in the model zoo for the source federation, and 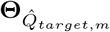 are the corresponding statistics for the target unseen site.

The chosen model can then either be used directly or adapted to the target site using the final part of the unified framework.

### Model Adaptation to an Unseen Site

The method for Model Adaptation to an Unseen Site largely follows the previously presented conference proceedings [33], but here is adjusted for application to 3D. We assume that the federation was trained previously and we no longer have access to the source data that was used to train the models, whether due to lack of data sharing agreements or the difficulties of storing data for long periods. We thus assume that we only have access to the source model trained on the data federation, with model parameters 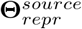 and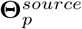, and the corresponding reference statistics 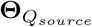. These do not require normalisation as we are using the features from a single model. For the target site, we have access to the data samples ***X***^*target*^ (*D*_*target*_ = *{****X***^*target*^*}*) and no labels are required.

The target model is initialised using the source trained weights, and the model training involves finetuning the feature extractor (following the naming conventions in Fig. 1), Θ_*repr*_, to align the feature distributions of the source and target models. Thus, the label predictor is frozen and the label predictor model weights remain fixed as 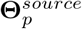. The finetuning is achieved by calculating the current features for the target site 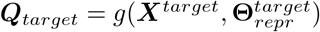 and estimating the Gaussian parameters, using the same subset of features *N*_*index*_ as used to train the source model:

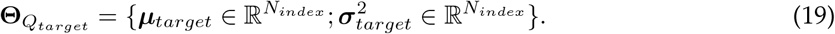

The loss function to finetune the model is then simply the Bhattacharyya distance between the source and target distributions:

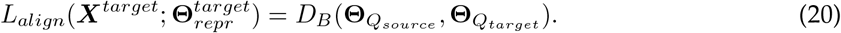

Through minimising this loss function we finetune the feature extractor to match the distribution, meaning that the source label predictor can now be used to produce model predictions such that 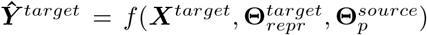, where *f* is the function mapping between the input images and target labels. This, therefore, ensures that, given data from the source model training federation or the target domain with the same feature embedding, the same label prediction is given across sites, and a new site can benefit from the knowledge of the data federation without requiring the data to still be available.

### Implementation Details

All comparison methods used the same network architecture, with features always extracted in the penultimate layer, before the activation function. Where necessary for adversarial methods, a discriminator was at the penultimate convolutional layer, after the last skip connection, following [36]. Training was completed on an A10 GPU using PyTorch 1.12.0. For the federated learning, a learning rate of 1 *×* 10^*−*4^ was used with an AdamW optimiser, and for the model adaptation, a learning rate of 1 *×* 10^*−*6^ was used with an AdamW optimiser.

## Acknowledgements

A.I.L.N. and N.K.D. are supported by the Bill and Melinda Gates Foundation. MJ is supported by the National Institute for Health Research, Oxford Biomedical Research Centre, and this research was funded by the Wellcome Trust [215573/Z/19/Z]. WIN is supported by core funding from the Wellcome Trust [203139/Z/16/Z]

## Footnotes

1 https://fsl.fmrib.ox.ac.uk/fsl/fslwiki/fsl_anat

