## Supplementary Material for "UniFed: A unified deep learning framework for segmentation of partially labelled, distributed neuroimaging data" (from another version of this preprint: version 1, 2024-02-06)

#### 1 Additional Dataset Details

Table 1 shows the number of samples for each site which passed the preprocessing pipelines, and the train test splits. Figure 1 then shows the confusion matrix from training on each site separately in turn and testing on all remaining sites, assuming all sites are fully supervised. It can clearly be seen that there is a high degree of domain shift present between some sites from the drop in test performance. The average performance for training and testing on the same individual site was $55.04 \pm 24.60$, clearly showing many sites had insufficient data to train a model directly and demonstrating the need to be able to pool data across sites.

| Site | Train | Test | Total |
|---|---|---|---|
| CMU | 21 | 6 | 27 |
| Caltech | 30 | 8 | 38 |
| KKI | 32 | 9 | 41 |
| Leuven | 42 | 8 | 50 |
| MaxMun | 36 | 10 | 46 |
| NYU | 144 | 37 | 181 |
| OHSU | 22 | 6 | 27 |
| Olin | 28 | 8 | 36 |
| Pitt | 44 | 11 | 55 |
| SBL | 24 | 6 | 30 |
| SDSU | 27 | 8 | 35 |
| Stanford | 31 | 8 | 39 |
| Trinity | 37 | 10 | 47 |
| UM | 107 | 28 | 135 |
| Yale | 44 | 12 | 56 |
| UCLA | 78 | 20 | 98 |

Table 1: Number of samples by site for ABIDE dataset post FSL Anat preprocessing

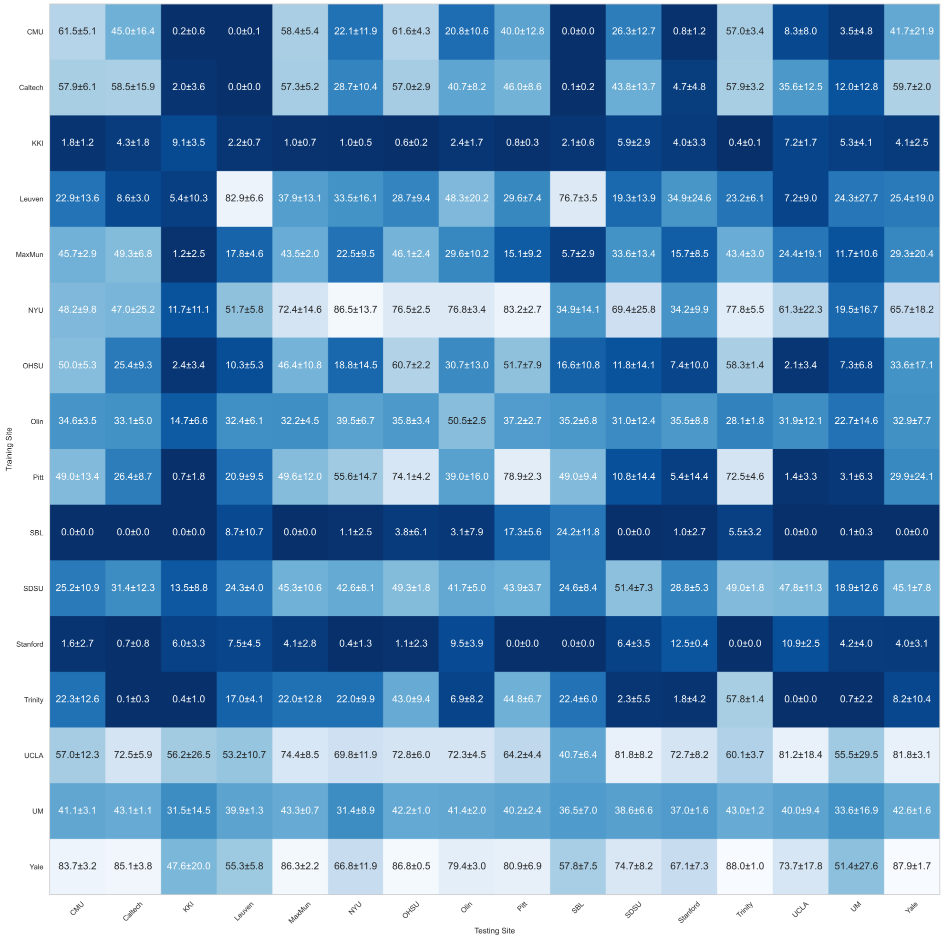


Figure 1: Confusion matrix from training on each site separately, full supervised.

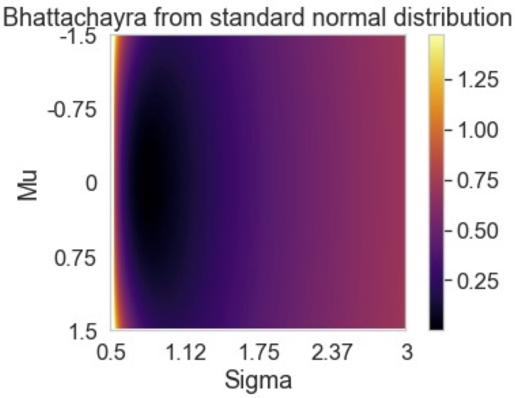


Figure 2: Bhattacharyya Distance for two GMMs: increasing loss as the second Gaussian increases in difference from the original reference Gaussian with $\mu = 0$ and $\sigma = 1$.

#### 2 Extension to Gaussian Mixture Models

The Bhattacharyya Distance is a general distance between two distributions, which can be used to measure the degree of similarity between two probability distributions. For two probability distributions P and Q on the same domain, the Bhattacharyya distance is defined as:

$$D_B(P,Q) = -ln(BC(P,Q)) \tag{1}$$

where

$$BC(P,Q) = \sum \sqrt{P(x)Q(x)} \tag{2}$$

is the Bhattacharyya coefficient for discrete probability distributions or

$$BC(p,q) = \int_x \sqrt{p(x)q(x)} dx. \tag{3}$$

for two continuous probability distributions $P(dx) \sim p(x)dx$ and $Q(dx) \sim q(x)dx$.

Thus if $p(x) = \mathcal{N}(\mu_p, \sigma_p)$ and $q(x) = \mathcal{N}(\mu_q, \sigma_q)$ then:

$$D_B(P,Q) = -ln\Big(\int_x \sqrt{p(x)q(x)} dx\Big) \tag{4}$$

$$D_B(P,Q) = -ln\Big(\int_x \frac{1}{\sqrt{2\pi}\sigma_p}\frac{1}{\sqrt{2\pi}\sigma_q} \exp\Big(-\frac{1}{2}(\frac{x-\mu_p}{\sigma_p})^2\Big) \exp\Big(-\frac{1}{2}(\frac{x-\mu_q}{\sigma_q})^2\Big)\Big) dx \tag{5}$$

and thus it can be shown that:

$$D_B(P,Q) = \frac{1}{4}\frac{(\mu_p - \mu_q)^2}{\sigma_p^2 + \sigma_q^2} + \frac{1}{2}\ln\Big(\frac{\sigma_p^2 + \sigma_q^2}{2\sigma_p\sigma_q}\Big). \tag{6}$$

This distance has several desirable properties for training neural networks. Firstly, it is zero when the two distributions are identical. Secondly, it is clearly differentiable. Figure 2 shows the Bhattacharyya Distance as the second distribution is increasingly removed from the standard normal distribution reference. The loss is clearly smoothly varying and also penalises the collapse of distribution - as $\sigma$ tends to 0, the loss rapidly increases in magnitude. It can also be observed that in the case where the $\sigma$ values are equal, the Bhattacharyya distance becomes the Mahalanobis distance between the two distributions. Thus, the Bhattacharyya distance is clearly the better choice, as it characterises differences between the distributions both in terms of the mean and also the standard deviation.

However, in this work we wish to consider the similarity between two Gaussian mixture models (GMM) with an arbitrary number of components. Note that for the Bhattacharyya distance, the two distributions being considered must be of the same type, and thus the reference and target GMM distributions must contain the same number of components. If we consider two GMMs defined as:

$$p(x) \sim \sum_{k=1}^{M} \pi_{pk} \mathcal{N}(x; \mu_{pk}, \sigma^2_{pk}) \tag{7}$$

and similarly for $q(x)$, where k is the number of components in the GMM and $\pi_p$ are the mixing weights, then if we calculated $D_B$ for the simplest case when $k = 2$:

$$D_B = -ln\Big(\int_x \sqrt{PQ}\Big)\Big)dx \tag{8}$$

where

$$P = \pi_p \frac{1}{\sqrt{2\pi}\sigma_{p1}} \exp\Big(-\frac{1}{2}(\frac{x-\mu_{p1}}{\sigma_{p1}})^2\Big) + (1-\pi_p)\frac{1}{\sqrt{2\pi}\sigma_{p2}} \exp\Big(-\frac{1}{2}(\frac{x-\mu_{p2}}{\sigma_{p2}})^2\Big) \tag{9}$$

and

$$Q = \pi_q \frac{1}{\sqrt{2\pi}\sigma_{q1}} \exp\Big(-\frac{1}{2}(\frac{x-\mu_{q1}}{\sigma_{q1}})^2\Big) + (1-\pi_q)\frac{1}{\sqrt{2\pi}\sigma_{q2}} \exp\Big(-\frac{1}{2}(\frac{x-\mu_{q2}}{\sigma_{q2}})^2\Big). \tag{10}$$

This integral is thus the square root of a sum of exponential functions and, thus, has no closed form solution.

Thus, we utilise the approximation:

$$D_{GMM}(P,Q) = \sum_{k=1}^{M} \pi_{pk}\pi_{qk}\Big(\frac{1}{4}\frac{(\mu_{pk}-\mu_{qk})^2}{\sigma^2_{pk}+\sigma^2_{qk}} + \frac{1}{2}\ln\Big(\frac{\sigma^2_{pk}+\sigma^2_{qk}}{2\sigma_{pk}\sigma_{qk}}\Big)\Big) \tag{11}$$

and clearly when P=Q $D_{GMM} = 0$.

### 3 Implementation Details

We here detail the implementation details for the comparison methods used in this work for each task in turn. As far as possible we used the recommended settings from the relevant papers, and kept the other implementation aspects as close to our proposed approach as possible. All models were trained with 5 fold cross validation, and the best fold used for test evaluation.

#### 3.1 Sites Labelled

**Centralised Data:** Learning rate = $1 \times 10^{-4}$ with AdamW optimiser. Trained with early stopping with a patience of 25 epochs.
**FedAvg:** Learning rate = $1 \times 10^{-4}$ with AdamW optimiser. Trained with early stopping with a patience of 25 epochs at local sites.
**FedProx:** Learning rate = $1 \times 10^{-4}$ with AdamW optimiser. Trained with early stopping with a patience of 25 epochs at local sites. Proximal term weight $\mu = 1 \times 10^{-4}$, chosen empirically.
**FedPerl:** Learning rate = $1 \times 10^{-4}$ with AdamW optimiser. Trained with early stopping with a patience of 25 epochs at local sites. Implemented using the source code available at https://github.com/tbdair/FedPerlV1.0. Empirically, T, the number of sites in the committee was set to 3, but similar performance was seen across a large range of T values.
**FSSL-DP:** Learning rate = $1 \times 10^{-4}$ with AdamW optimiser. Trained with early stopping with a patience of 25 epochs at local sites. $\lambda$, the uncertainty threshold was set to 0.9.
**FADA [1]:** The model was trained at the local site until convergence with a patience of 10 epochs. The model was trained with a learning rate $1 \times 10^{-4}$ with AdamW optimiser. Implemented using the supplied code: https://drive.google.com/file/d/1OekTpqB6qLfjlE2XUjQPm3F110KDMFc0/view.
**FedHarmony [2]:** Implemented using the supplied code: https://github.com/nkdinsdale/FedHarmony. The model was trained at the local site until convergence with a patience of 10 epochs. The model was trained with a learning rate $1 \times 10^{-4}$ with AdamW optimiser. The recommended hyperparameters were used apart from for the proximal term which was tuned to the task: $\mu = 0.0001, \alpha = 1, \beta = 100$.

##### 3.2 Percentage Labelled

As above plus:
**Omnisupervised:** Learning rate = $1 \times 10^{-4}$ with AdamW optimiser. Trained with early stopping with a patience of 25 epochs at local sites. Followed the approach in [3], and dropout value $p = 0.5$.

##### 3.3 Model Adaptation

As above plus:
**Target finetune:** Learning rate = $1 \times 10^{-5}$ with AdamW optimiser. Trained with early stopping with a patience of 25 epochs at local sites.
**Minimise Entropy:** Vanilla entropy minimisation. The model was trained with a learning rate $1 \times 10^{-5}$ with AdamW optimiser. The model was trained at the local site until convergence with a patience of 25 epochs.
**AdaEnt [4]:** Implemented using the supplied code: `https://github.com/mathilde-b/SFDA`. The hyperparameter weighting loss function contributions was set to 100 following the paper. The model was trained with a learning rate $1 \times 10^{-5}$ with AdamW optimiser. The model was trained at the local site until convergence with a patience of 25 epochs.
**AdaMI [5]:** Implemented using the supplied code: `https://github.com/mathilde-b/SFDA`. The hyperparameter weighting loss function contributions was set to 100 following the paper. The tissue prior was estimated by averaging the source site labels and then the tissue ratio was used to create a slice depth dependent tissue prior. The model was trained with a learning rate $1 \times 10^{-5}$ with AdamW optimiser. The model was trained at the local site until convergence with a patience of 25 epochs.
**MMD** The model was trained with a learning rate $1 \times 10^{-5}$ with AdamW optimiser. Trained with early stopping with a patience of 25 epochs. MMD was implemented instead of DeepCORAL [6], as DeepCORAL could not be implemented due to memory constraints.

##### 3.4 Model Architecture

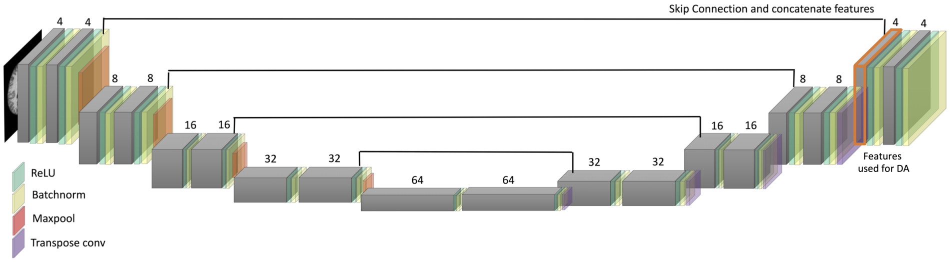


Figure 3: UNet model architecture.

A 3D UNet was trained on the MRI volumes, with 4 max pooling layers and 2 convolutional blocks per depth. An initial feature number of 4 was used. The highlighted features represent the features modelled by the Gaussian embedding, after the last skip connection. Again, the key architecture choice is that the embeddings are returned before the ReLU function, so that they are better represented by a GMM and do not have a sharp cut off.

### 4 Training Graphs

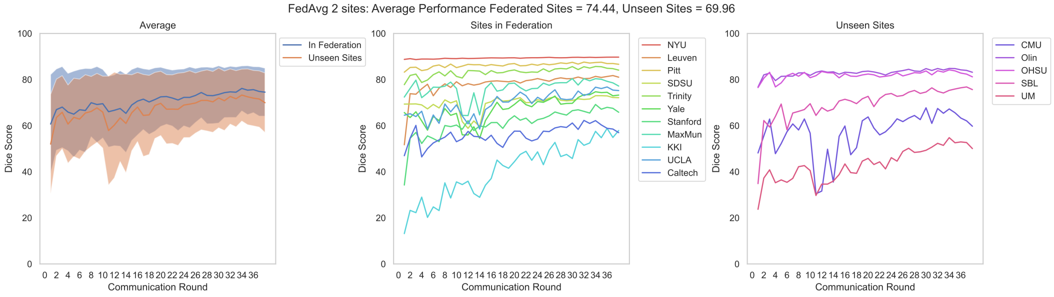


(a) FedAvg Training Graphs 2 Sites Supervised

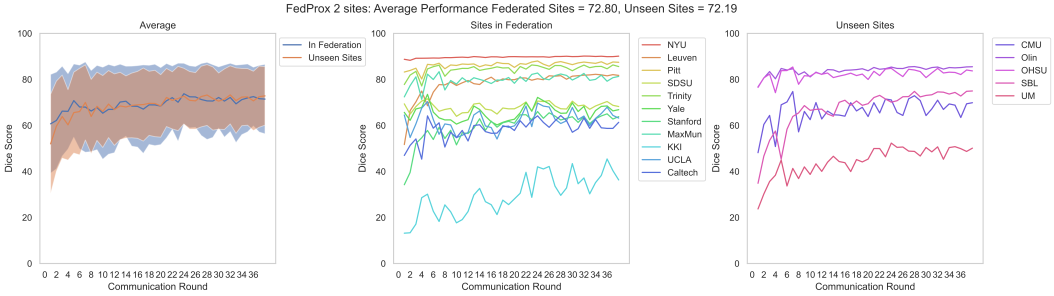


(b) FedProx Training Graphs 2 Sites Supervised

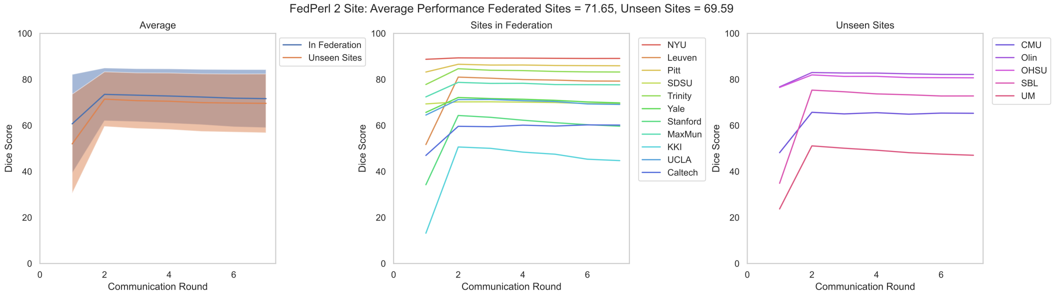


(c) FedPerl Training Graphs 2 Sites Supervised

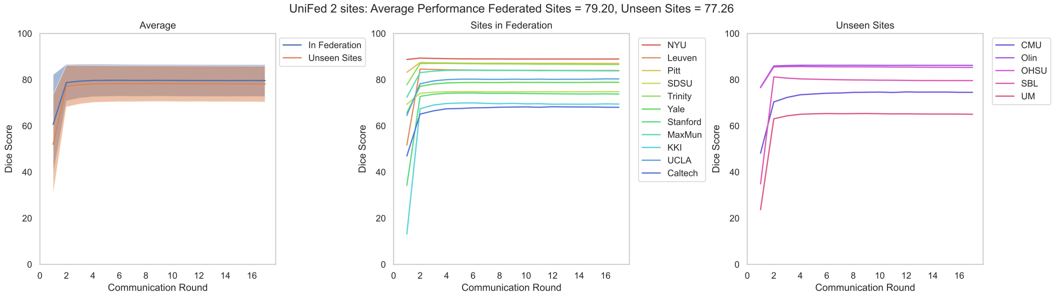


(d) UniFed Training Graphs 2 Sites Supervised

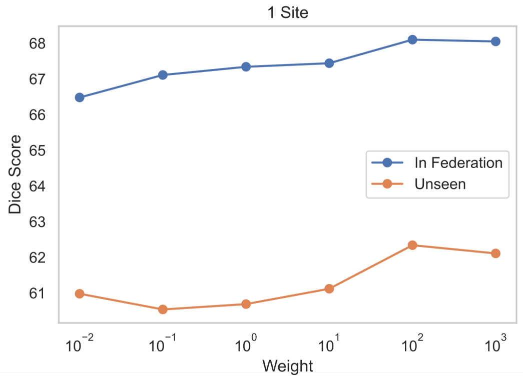


(a) 1 Site

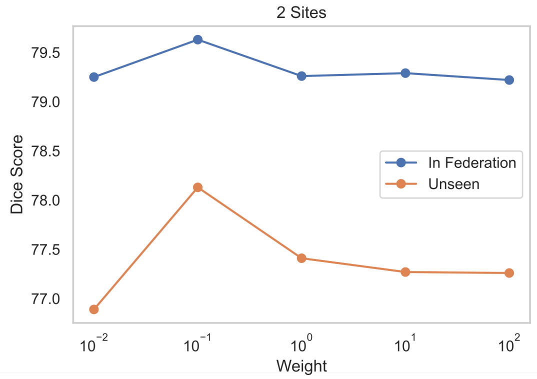


(b) 2 Sites

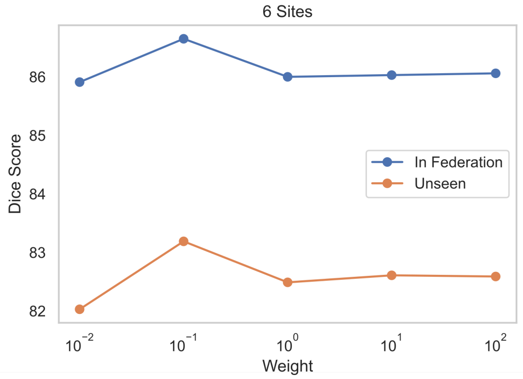


(c) 6 Sites

Figure 5: Hyper parameter sensitivity results

| Method | 2 Sites | | 6 Sites | |
|---|---|---|---|---|
| | Mean Federation | Mean Unseen | Mean Federation | Mean Unseen |
| FedAvg [7] | 85.11 | 72.80 | 86.11 | 84.89 |
| FSSL-DP [8] | 57.33 | 46.70 | 85.69 | 83.65 |
| FADA [1] | 84.56 | 74.50 | 88.90 | 87.69 |
| FedHarmony [9] | 84.46 | 73.04 | 89.62 | 87.54 |
| UniFed | **85.14** | **76.12** | **92.20** | **89.32** |

Table 2: Results for alternative sites split

### 5 Additional Results

#### 5.1 Sensitivity to hyper parameter

Our approach only introduces a single hyper parameter, the weighting value between the main task loss and the Bhattacharyya distance. Figure 5 shows the effect of varying the hyper parameter for 1, 2 and 6 sites supervised. It can be seen that the performance is robust to the choice of the hyper parameter.

#### 5.2 Effect of random split

We only completed the main experiments for a single random split of sites into the federation and unseen sites. We repeated the experiment for a different random split for 2 and 6 sites supervised, with Yale as the reference site. The replication was limited to this due to the high computation times for experiments. The results can be seen in Table 2, where it can be seen that the pattern of results was the same, with UniFed outperforming existing methods.
